# Cellular orientational fluctuations, rotational diffusion and nematic order under periodic driving

**DOI:** 10.1101/2022.04.30.490160

**Authors:** Avraham Moriel, Ariel Livne, Eran Bouchbinder

## Abstract

The ability of living cells to sense the physical properties of their microenvironment and to respond to dynamic forces acting on them plays a central role in regulating their structure, function and fate. Of particular importance is the cellular sensitivity and response to periodic driving forces in noisy environments, encountered in vital physiological conditions such as heart beating, blood vessels pulsation and breathing. Here, we first test and validate two predictions of a mean-field theory of cellular reorientation under periodic driving, which combines the minimization of cellular anisotropic elastic energy with active remodeling forces. We then extend the mean-field theory to include uncorrelated, additive nonequilibrium fluctuations, and show that the theory quantitatively agrees with the experimentally observed stationary probability distributions of the cell body orientation, under a range of biaxial periodic driving forces. The fluctuations theory allows to extract the dimensionless active noise amplitude of various cell types, and consequently their rotational diffusion coefficient. We then focus on intra-cellular nematic order, i.e. on orientational fluctuations of actin stress fibers around the cell body orientation, and show experimentally that intra-cellular nematic order increases with both the magnitude of the driving forces and the biaxiality strain ratio. These results are semi-quantitatively explained by applying the same cell body fluctuations theory to orientationally correlated actin stress fiber domains. The implications of these findings, which make the quantitative analysis of cell mechanosensitivity more accessible, are discussed.

## INTRODUCTION

Cells in our body exhibit a high degree of spatial organization in complex, dynamic and noisy biophysical environments. The highly-ordered spatial cellular organization is essential for the biological functions of single cells, as well as for the tissues and organs that they form, and even for their fate. In many cases, cells are exposed to high-frequency periodic driving forces in addition to intrinsic and extrinsic noise, e.g. in vital physiological conditions such as those encountered in cardiovascular tissues in hemodynamic environments, in the lungs under breathing motion and in cardiac tissue under rhythmic heart beating. The noise under such conditions is clearly of nonequilibrium and active nature. The absence of proper cellular organization may lead to dysfunction, pathological conditions and diseases such as hypertrophic cardiomyopathy [1–4]. Furthermore, it is now recognized that retaining cellular organization is of crucial importance for tissue engineering and cardiovascular regenerative medicine [5–11].

A major challenge is to quantitatively understand the relations between the mechanical forces applied to the cellular environment, the fluctuations cells experience and exhibit, their relevant cellular properties and the resulting dynamic response. The latter requires cells to “sense” some physical properties of their environment and to actively tune their subsequent dynamic response, which is therefore termed mechanosensitive.

Well-controlled laboratory experiments, in which living cells adhere to a deformable substrate that is exposed to periodic driving forces [12–26], provide an excellent test bed for probing basic aspects of cellular mechanosensitivity in driven systems and for extracting quantitative information about the accompanying active fluctuations. A cartoon of such experiments is shown in Fig. 1a, presenting an ensemble of non-interacting cells (ellipses) adhering to a deformable 2D substrate (of elastic stiffness *E*_sub_), which mimics the extracellular matrix in physiological conditions. The substrate is exposed to cyclic stretching of amplitude *ϵ* and frequency *f* in the horizonal direction (here the stretch is extensional) and to cyclic stretching of amplitude *−r ϵ* (compressional) and the same frequency *f* in the perpendicular direction.

**FIG. 1.**
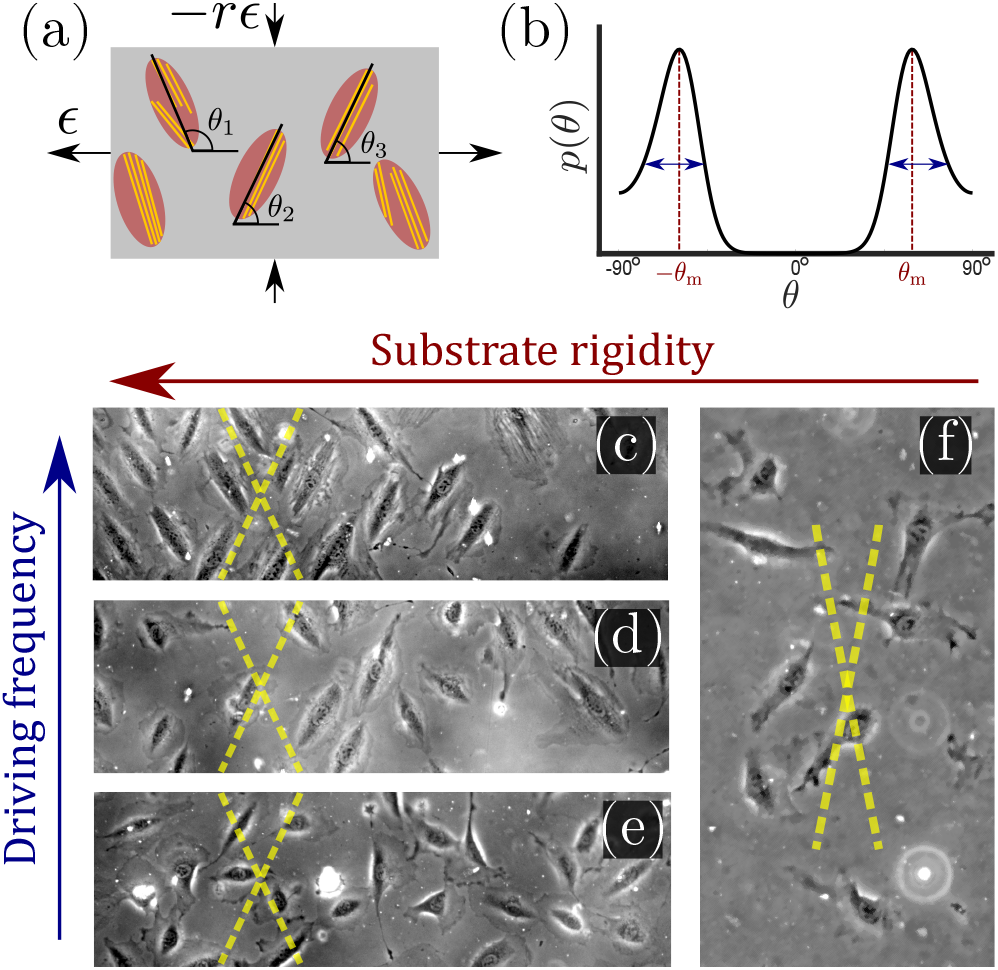
(a) A cartoon of the setup used in cyclic stretching experiments, where non-interacting adherent cells (ellipses) are exposed to periodic driving forces applied to the underlying substrate (grey rectangle). The parameters of the driving force — the strain amplitude *ϵ* and the biaxiality strain ratio *r* — are explicitly indicated (see text for additional discussion). The other driving parameter — the driving frequency *f* defined in Eq. (1), is not shown. The cell body orientation *θ*_*i*_ (relative to the principal strain direction), of some of the cells, is marked. Finally, actin stress fibers, whose orientation does not necessarily coincide with the cell body orientation, are also sketched (yellow lines). (b) A sketch of a typical bimodal distribution *p*(*θ*), corresponding to the stationary cell body orientations of an ensemble of cells as those illustrated in panel (a). It features peaks at two mirror-image angles *±θ*_m_ (marked in subsequent panels by yellow dashed lines) and a finite characteristic width, illustrated by the double-headed arrows. (c) A phase-contrast image of rat embryo fibroblast (REF-52) cells on a fibronectin-coated PDMS substrate (*E*_sub_ ≃1 MPa), after 11 h of periodic stretching (*ϵ* = 0.1, *r* = 0.25, *f* = 1.2 Hz) demonstrates that originally randomly oriented cells (not shown) are aligned in one of the two mirror-image orientations, marked by the yellow dashed lines [26]. In this experiment, the conditions *f ≫τ* ^*−*1^ *∼* 0.1 Hz (where *τ* is an intrinsic cellular timescale, see text for additional details) and *E*_sub_ ≫ *E*_eff_ are met. (d) The same as panel (c), but with *f* = 0.12 Hz, which corresponds to *f ≃ τ* ^*−*1^. In this case, not all cells are aligned in one of two mirror-image orientations (partial reorientation). (e) The same as panels (c) and (d), but with *f* = 0.008 Hz, which corresponds to *f ≪ τ* ^*−*1^. In this case, cells are not aligned at all in special directions. (f) The same as panel (c), but with *E* _sub_ ≃5 kPa, which corresponds to *E*_sub_ ≲*E*_eff_ (here = 0.15). In this case, cells are not aligned in special directions.

Upon applying the biaxial cyclic stretching for a long time, each cell settles into a well-defined orientation/angle *θ*_*i*_ relative to the principal direction. Each cell also features intra-cellular cytoskeletal organization of its actin stress fibers (yellow lines), whose orientation does not necessarily coincide with the cell body orientation *θ*_*i*_. The orientations {*θ*_*i*_} are typically clustered around two mirror-image angles, but generically correspond to a continuous distribution *p*(*θ*). Such a characteristic bimodal distribution is sketched in Fig. 1b, where the mirror-image angles — denoted by *±θ*_m_ — correspond to the symmetric peaks of the distribution. Each peak also features a finite width, marked by the double-headed arrows. Many cell types are known to undergo reorientation dynamics in response to such periodic driving [12–26], for sufficiently rigid substrates and high frequencies *f*, as demonstrated in Fig. 1c for rat embryo fibroblast (REF-52) cells.

In mathematical terms, in these cyclic stretching experiments the following two-dimensional, time-dependent strain tensor

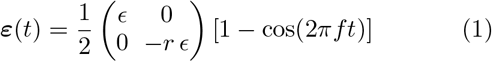

is applied to the substrate to which the cells adhere. As explained above, this periodic driving is characterized by three parameters: the strain *ϵ* in the principal direction, the strain *−r ϵ* in the perpendicular direction — expressed here in terms of the biaxiality strain ratio *r* —, and the frequency *f*. The application of this periodic driving triggers a mechanosensitive cellular response in the form of time-dependent reorientation dynamics, which for long times result in stationary orientational distributions *p*(*θ*) of the type illustrated in Fig. 1b. A major challenge is to understand in quantitative terms these orientational dynamics and the emerging statistical distributions, along with their dependence on both the external conditions and internal cellular properties.

The experimental observations described above have triggered quite a lot of theoretical work, for example [26– 34]. A mean-field theory, to be briefly reviewed below, quantitatively predicted the entire reorientation dynamics of single cells, from an initial random orientation to a well-defined final orientation [26]. The very same single-cell theory, which combines the minimization of anisotropic cellular elastic energy with active remodeling forces, has been used to explain orientational order in vascular networks formed by a coculture of fibroblasts and endothelial cells embedded in three-dimensional biomaterials under periodic driving [9]. Very recently, it has also been used to account for the orientational order of Caco-2 cells inside confluent epithelial layers under periodic driving [35]. These latter works demonstrate that mechanosensitivity and the accompanying orientational order at the cellular level can significantly affect tissue structures and functions at higher levels of organization. Here, our goal is to understand the stationary distributions *p*(*θ*) of cellular orientation under periodic driving, as well as the accompanying cytoskeletal organization of cells, most notably the orientation of actin stress fibers (SFs). The latter allows to quantify the degree of intra-cellular nematic order under periodic driving. To achieve these goals, we first test and validate two predictions of the a mean-field theory of cellular reorientation under periodic driving. We then extend the theory to include uncorrelated, additive nonequilibrium fluctuations. Using experimental data for various cell types, we show that the fluctuations theory quantitatively agrees with the experimentally observed stationary probability distributions of cell body orientation, under a range of biaxial periodic driving forces. These results demonstrate that the anisotropic elastic energy of cells, in the highfrequency regime, plays a decisive role not just in their reorientation dynamics [26], but also in their orientational fluctuations, and support the validity of the uncorrelated, additive noise description.

The cell body analysis allows to extract the dimensionless amplitude of nonequilibrium orientational noise, or alternatively the rotational diffusion coefficient of cells. These cellular properties, when compared across various cell types, reveal interesting similarities and differences. We then shift our focus to the internal cytoskeletal organization of cells, as manifested by intra-cellular nematic order [36–39], i.e. by orientational fluctuations of actin SFs around the cell body orientation. We experimentally show that intra-cellular nematic order increases with both the magnitude *ϵ* of the driving forces and with the biaxiality strain ratio *r*. These results are semiquantitatively explained by applying the same cell body fluctuations theory to orientationally correlated stress fiber domains. Finally, the implications of our findings, as well as future investigation directions, are discussed.

### CELLULAR REORIENTATION UNDER PERIODIC DRIVING AND THE SINGLE-CELL MEAN-FIELD THEORY

Our first goal is to briefly review the deterministic (mean-field) single-cell theory of cellular reorientation under periodic driving, originally introduced and validated in [26], and to further substantiate it as a preliminary step for discussing orientational fluctuations. The theory considers a cell that adheres to a deformable substrate, which is subjected to the two-dimensional strain tensor of Eq. (1), and that is oriented at time *t* at an angle *θ*(*t*) relative to the largest principal strain direction. It then derives the time-averaged linear elastic energy in the cell in form 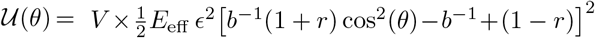, where *V* is the effective cell volume, *E*_eff_ is the effective cell stiffness and *b* is a dimensionless parameter that characterizes the elastic anisotropy of the cell. The elastic energy 𝒰(*θ*) has been averaged over a timescale *τ* significantly longer than *f* ^*−*1^, such that *f* does not appear in the problem any more (the physical meaning of the *τ* and the condition *τ* »*f* ^*−*1^ will be further discussed below). The expression for 𝒰(*θ*) also assumes that all of the substrate deformation is transferred to the cell, i.e. that *E*_eff_ is significantly smaller than the substrate’s stiffness/rigidity *E*_sub_.

The time evolution of *θ*(*t*) has been suggested to follow the deterministic relaxational dynamics *dθ*(*t*)/*dt* = − *η*^−1^*d* 𝒰(*θ*)/*dθ* ≡−*τ*^−1^*dū*(*θ*)/*dθ*,, where *η* is an effective orientational viscous-like parameter, 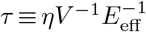 is an intrinsic cellular timescale (corresponding to active remodelling of actin structures and focal adhesions [40–42]) and

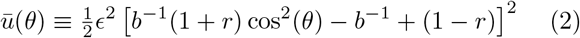

is the dimensionless elastic energy. These single-cell dynamics have been shown to be in remarkable quantitative agreement with the experimentally measured cell body orientation *θ*(*t*) of REF-52, for a wide range of experimental conditions. Moreover, it has been shown that for the fibroblasts used therein, *τ* = 6.6 *±* 0.4 sec and *b* = 1.13*±* 0.04 [26].

The value *τ* = 6.6 sec implies that the theory described above is valid for frequencies *f* » *τ* ^*−*1^ ∼0.1 Hz, and consequently that *E*_eff_ and *b* characterize the elastic cellular response in this high-frequency limit. This timescale separation predicts that in the opposite limit, *f* ≪ *τ* ^*−*1^ ∼0.1 Hz, the phenomenon disappears (at least for the used fibroblasts) since in this limit the cell has enough time to reduce its stored elastic energy by reducing its elastic stiffness (through active remodeling), not by reorientation. This prediction is verified in Fig. 1c-e, where *f* is varied from *f* » *τ* ^*−*1^ (panel (c)) to *f* « *τ* ^*−*1^ (panel (e)). Similar results for human mesenchymal stem cells have been very recently reported in [23], see in particular Fig. S9 in the Supporting Information file therein.

We can likewise test the assumed stiffness scale separation, which predicts that for *E*_sub_ »*E*_eff_ complete cellular reorientation would be observed, while for *E*_sub_ «*E*_eff_ the phenomenon disappears. In [26], complete cellular reorientation has been observed down to *E*_sub_ ≃20 kPa. In Fig. 1f, we present results for *E*_sub_≃5 kPa, where complete cellular reorientation is not observed. As such, the results support the prediction and indicate that the effective cell stiffness — which is a “composite” cellular property that is difficult to probe directly — is of the order of 10 kPa for REF-52. In fact, the described procedure offers a method for estimating the effective stiffness of cells in the high-frequency limit.

### FLUCTUATIONS IN CELLULAR ORIENTATION

The cellular reorientation process discussed above is robust, yet it is subjected to various sources of noise and fluctuations. The success of the single-cell theory in quantitatively predicting the cellular reorientation process at the mean-field level opens the way for developing a quantitative understanding of the fluctuations that accompany cellular orientation under driven conditions, which are of intrinsically nonequilibrium, active nature. While active fluctuations are found everywhere in living systems, their accurate quantification and understanding is generally difficult and nontrivial. Consequently, we focus next on the statistical properties of cellular orientation of ensembles of cells under periodic driving of the type presented in Eq. (1).

To this aim, we consider nonlinear Langevin dynamics of the form (as was also invoked, e.g., in [18, 35, 43])

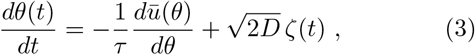

where *D* is an effective rotational diffusion coefficient (of inverse time dimension) and *ζ*(*t*) is assumed to be an uncorrelated noise with zero mean and unit variance, i.e. ⟨ *ζ*(*t*)⟩ = 0 and ⟨ *ζ*(*t*)*ζ*(*t*^*I*^) = *δ*(*t− t*^*I*^)⟩. Equation (3) reduces to the deterministic mean-field theory when averaged over the noise and contains a single new dimensionless parameter, *τ D* (recall that *τ* is determined from the reorientational dynamics of single cells in the framework of the mean-field theory). Our main goal is to understand whether the inclusion of uncorrelated additive orientational fluctuations, which are related to active/nonequilibrium noise (that cannot possibly be inherited from ordinary thermal fluctuations), quantitatively predicts the stationary probability distribution function *p*(*θ*) of cellular orientation in the long-time limit, and if so, to understand the properties of the orientational noise amplitude.

A standard procedure that invokes the Fokker-Planck equation, corresponding to the Langevin dynamics in Eq. (3), leads to the following stationary probability distribution function

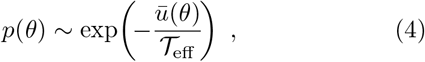

where 𝒯_eff_ ≡*τ D* is a dimensionless effective temperature that replaces the (properly normalized) ordinary temperature in thermal equilibrium and *θ* represents the long-time (steady state) cellular orientation. Combining Eq. (2) with Eq. (4), we obtain the following prediction for the most probable orientation *θ*_m_ (corresponding to the minimum of *ū*(*θ*))

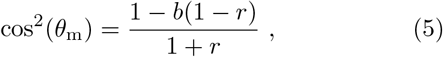

valid for 1 *−*1*/b* ≤ *r* ≤1*/*(*b −*1), which we focus on in this work. For 0 *<r<* 1−1*/b*, we predict *θ*_m_ = 90^*°*^. The former prediction has been extensively verified experimentally in [26], while the latter has been verified in [9]. In the present context, Eq. (4) implies that the location of the maximum of *p*(*θ*), *θ*_m_, uniquely determines the cellular parameter *b* (recall that the biaxiality ratio *r* is externally imposed).

To test the prediction of Eqs. (2)-(4), we consider first the experimental data of [18] for the steady-state cellular orientation of primary human umbilical cord fibroblasts, under the periodic driving of Eq. (1) (obtained under the conditions *f* » *τ* ^*−*1^ and *E*_sub_ » *E*_eff_). The reason we first focus on these experimental data is that they offer — to the best of our knowledge — the largest cell ensembles currently available, including a few hundreds cells per each experimental condition (ϵ*E, r*), going up to ∼ 1500 cells in the best case. For comparison, the experiments of [26], which where focused on single-cell reorientation *dynamics* rather than on *stationary orientational statistics*, featured 𝒪(100) cells per experimental condition.

In Fig. 2, we present the cellular orientation distributions *p*(*θ*) (discrete histograms, to be precise, see the circles) for primary human umbilical cord fibroblasts, digitized from Fig. 4 of [18], for various *ϵ, r* and *f* conditions (cf. Table I). The most probable orientations corresponding to these distributions have already been analyzed in [26] and shown to be in quantitative agreement with the prediction in Eq. (5). Consequently, primary human umbilical cord fibroblasts are expected to feature elastic anisotropy close to *b* = 1.13 *±* 0.04, which characterizes REF-52. The periodic driving frequency *f* is in the range of tens of mHz (see Table I), i.e. about an order of magnitude smaller than the lowest frequency for which REF-52 still exhibit complete reorientation (cf. Fig. 1d). This implies that *τ* for primary human umbilical cord fibroblasts is more than one order of magnitude larger than that of REF-52. This point will be further discussed below.

**FIG. 2.**
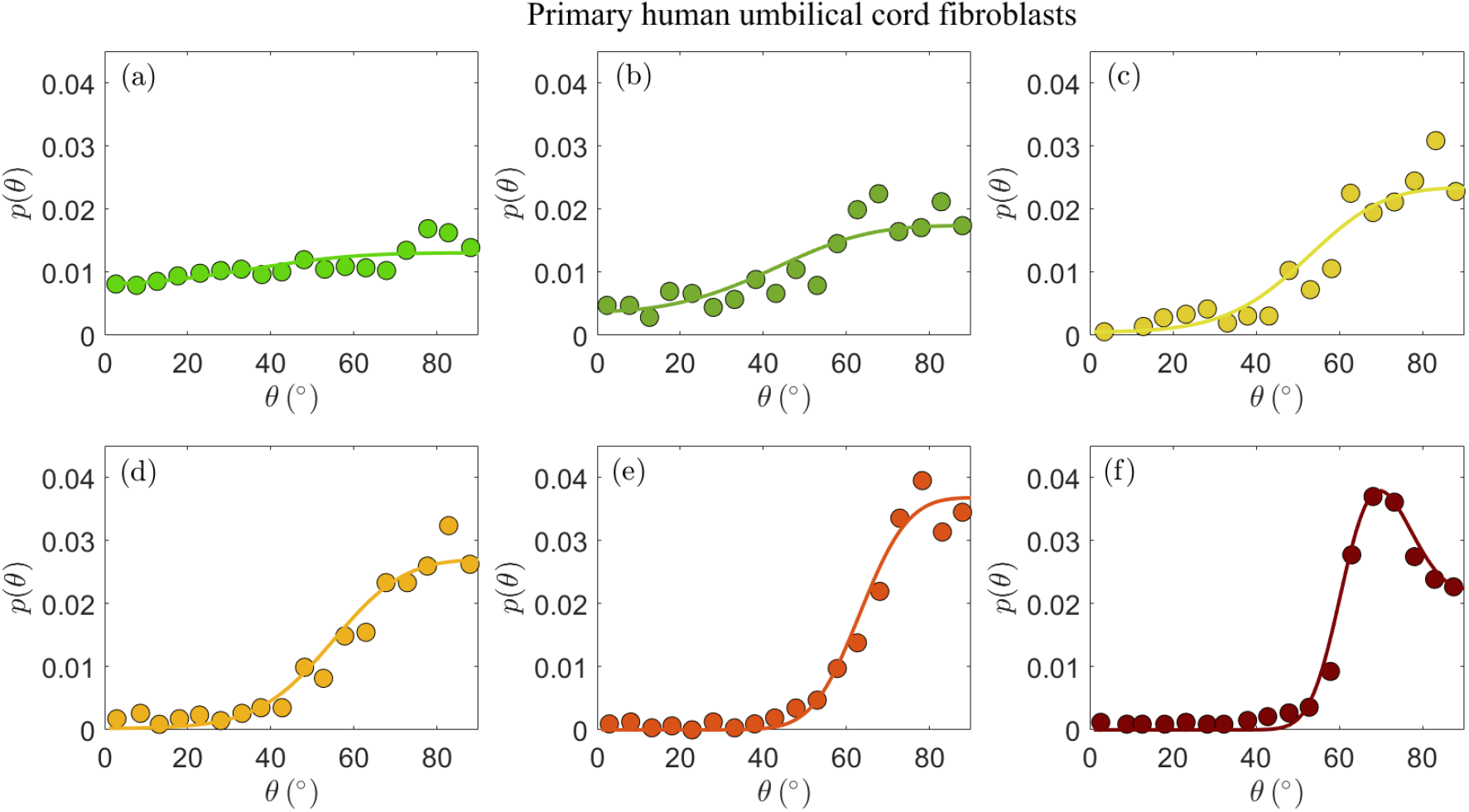
The stationary probability distribution function *p*(*θ*) (circles) of the orientation of primary human umbilical cord fibroblasts on a fibronectin-coated PDMS substrate (*E*_sub_ ≃50 kPa) after 16 h of periodic stretching [18]. Each panel corresponds to different imposed parameters *ϵ, r* and *f* in Eq. (1), as detailed in Table I (see text for the discussion of the *r* values). The circles correspond to the data appearing in Fig. 4 of [18]. The solid lines correspond to theoretical fits to Eq. (4) (together with Eq. (2)), obtained through the procedure detailed in the text. The values of the dimensionless cellular parameters *b* and *T*_eff_ appear in Table I and are discussed in the text.

**TABLE I.** The experimental parameters [18] and the extracted cellular parameters corresponding to panels (a)-(f) of Fig. 2, see extensive discussion in the text.

| panel | $\epsilon$ | $r$ | $f$ (mHz) | $b$ | $\mathcal{T}_{\text{eff}}$ |
| --- | --- | --- | --- | --- | --- |
| (a) | 4.9% | 0.10 | 52 | 1.20 | $2.5 \times 10^{-3}$ |
| (b) | 8.4% | 0.10 | 34 | 1.20 | $2.2 \times 10^{-3}$ |
| (c) | 11.8% | 0.10 | 25 | 1.20 | $1.8 \times 10^{-3}$ |
| (d) | 14.0% | 0.10 | 21 | 1.20 | $2.0 \times 10^{-3}$ |
| (e) | 32.0% | 0.12 | 9 | 1.16 | $3.4 \times 10^{-3}$ |
| (f) | 31.7% | 0.28 | 9 | 1.18 | $1.6 \times 10^{-3}$ |

The experimental data in Fig. 2a-d feature *ϵ* = 0.049 *−*0.140 and biaxiality ratios that are reported to be in the range *r* = 0.15 *±*0.05 [18]. We found that these data are consistent with *b* = 1.2 (i.e. close to the upper end of the *b* values of REF-52) and with *r* = 0.10 (i.e. in the lower end of the reported range); fixing *b* and *r* to these values, we fitted the experimental data in Fig. 2a-d to Eq. (4), together with Eq. (2), where 𝒯_eff_ is a free parameter in each dataset. The resulting fits are superimposed on Fig. 2a-d (solid lines) and the extracted values of 𝒯_eff_ are reported in Table I. The theoretical fits are in favorable agreement with the experimental data, and 𝒯_eff_ =(2.15 0.35) × 10^*−*3^ appears to be roughly independent of *ϵ* for this single *r* value.

The experimental data in Fig. 2a-d have been very recently analyzed in [43] using essentially the same theory (based on the previous work of some of us [26]). Though the analysis procedure and parameter values are slightly different (see [43] for details), favorable agreement with the theory has been also demonstrated, consistently with our findings. We next consider the data presented in Fig. 2e-f, which have not been analyzed in [43]. These two datasets have been obtained under significantly larger strains, *ϵ* ≃ 0.32 (cf. Table I), and two biaxiality ratios (*r* = 0.15 *±*0.05 in panel (e) and *r* = 0.29 *±*0.05 in panel (f)). The larger *ϵ* value may imply that the quadratic energy function of Eq. (2) requires higher order corrections in *ϵ* however, recent work has demonstrated that the steady state relation in Eq. (5), which corresponds to the minimum of *ū*(*θ*) in Eq. (2), remains valid for a very large class of nonlinear constitutive relations, characterised by orthotropic energy functionals [33]. Consequently, we analyze the experimental in Fig. 2e-f using the same theory discussed above.

Focusing first on Fig. 2e, we again constrain *r* in the range 0.10−0.20 and *b* in the range 1.16−1.2, and use the dimensionless noise amplitude 𝒯_eff_ as a free fitting parameter. The resulting theoretical fit, with *r* = 0.12, *b* = 1.16 And 𝒯_eff_ = 3.4 × 10^*−*3^ (cf. Table I) is superimposed on the experimental data (solid line), again demonstrating favorable agreement with the data. The extracted noise amplitude 𝒯_eff_ = 3.4 × 10^*−*3^ is somewhat larger than the approximately *ϵ*-independent value extracted for smaller*ϵ, T*_eff_ =(2.15 *±* 0.35)*×*10^*−*3^. At present, we cannot conclusively determine if the larger 𝒯_eff_ is a signature of a dependence on the strain *ϵ* or whether it is related to other effects.

The experimental dataset presented in Fig. 2f is in some sense the most interesting one, posing the most stringent test of the theoretical predictions. The reason for this is not just the fact that its biaxiality ratio is different, *r* = 0.29 *±*0.05, but mainly that it is the only dataset that features a clear maximum (attained at an orientation denoted by *θ*_m_). The latter is of importance because it requires *r* and *b*, which are anyway strongly constrained, to satisfy Eq. (5). This relation is indeed satisfied with *r* = 0.28 and *b* = 1.18, well within their narrow allowed ranges. With *r* = 0.28 and *b* = 1.18 fixed, we fit the theory to the experimental distribution *p*(*θ*) using a single free parameter 𝒯_eff_. The theoretical prediction (solid line) is in excellent agreement with the experimental data. The extracted noise amplitude 𝒯_eff_ = 1.6 × 10^*−*3^ is about half of that of the data in Fig. 2e, which feature a smaller *r* value. In the absence of a systematic variation of *r* over a broader range, we are currently unable to determine whether 𝒯_eff_ indeed reveals some robust *r* dependence for a fixed *ϵ* (note that some *r* dependence is encapsulated in the energy functional of Eq. (2)).

Overall, the comparison in Fig. 2 of the theoretical prediction of Eqs. (2) and (4) to the experimental data of primary human umbilical cord fibroblasts provides support for the cellular orientation fluctuations theory we consider here. In particular, it supports the existence of uncorrelated, additive nonequilibrium fluctuations (cf. Eq. (3)) — characterized by a dimensionless noise amplitude 𝒯_eff_ — and further strengthens the validity of the elastic anisotropic energy functional in Eq. (2), which has already been confirmed by dynamic reorientation measurements in [26]. Our next goal is to test whether these conclusions remain valid for other cell types, and if so, what can one learn from the extracted nonequilibrium noise amplitude *𝒯*_eff_.

To this aim, we consider *p*(*θ*) for human aortic endothelial cells, reported on in Fig. 5B of [19], which is presented here in Fig. 3a (digitized from the original figure. The same dataset has been analyzed in [32], using a somewhat related approach). This experiment has been performed under imposed strain of *ϵ* = 0.1 and biaxiality ratio of *r* = 0.34 [19]; since *p*(*θ*) in Fig. 3a features a clear maximum, Eq. (5) uniquely determines the cellular elastic anisotropy to be *b* = 1.23. This is, in itself, an important result as it shows that three different cell types feature very similar levels of elastic anisotropy quantified by *b*: REF-52 with *b* = 1.13 *±*0.04 [26], primary human umbilical cord fibroblasts with *b* = 1.18 *±*0.02 (cf. Fig. 2 and Table I) and human aortic endothelial cells with *b* = 1.23.

**FIG. 3.**
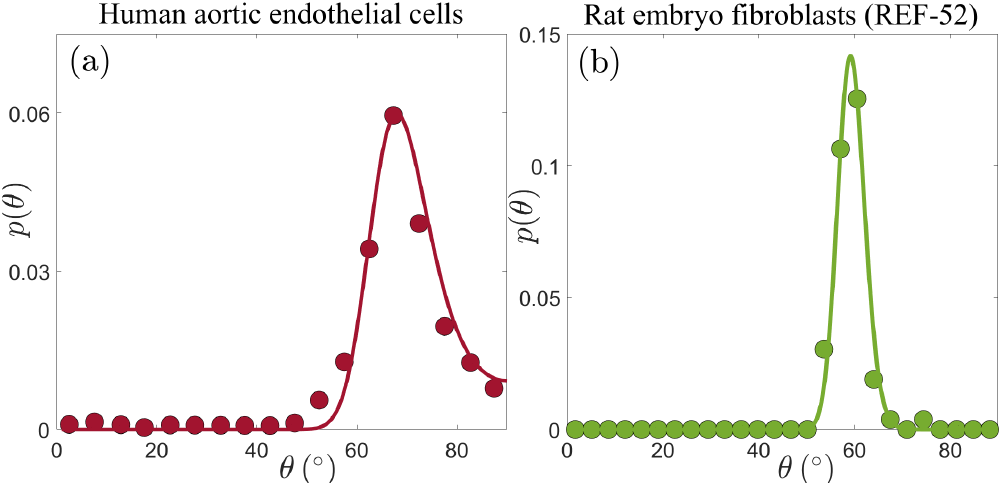
(a) *p*(*θ*) for human aortic endothelial cells (circles, extracted from Fig. 5B in [19], based on an ensemble of 77 − 144 cells), under periodic driving with *ϵ* = 0.1 and *r* = 0.34. The solid line corresponds to the theoretical prediction in Eq. (4) (together with Eq. (2)), with *b* = 1.23 and 𝒯_eff_ = 6.2 × 10^*−*5^ (see text for additional details). (b) *p*(*θ*) for REF-52 (circles, our experimental data, based on an ensemble of 76 cells), under periodic driving with *ϵ* = 0.104 and *r* = 0.46. The solid line corresponds to the theoretical prediction in Eq. (4) (together with Eq. (2)), with *b* = 1.15 and *T*_eff_ = 3.2 × 10^*−*5^ (see text for additional details).

With *ϵ* = 0.1, *r* = 0.34 and *b* = 1.23, we fit *p*(*θ*) in Fig. 3a to Eqs. (2) and (4) using a single free parameter 𝒯_eff_. The result is superimposed on the experimental data (solid line), revealing excellent agreement. The extracted dimensionless noise amplitude is 𝒯_eff_ = 6.2*×*10^*−*5^, which is a factor ∼35 smaller than the value extracted for primary human umbilical cord fibroblasts. What is the origin of this large difference in the dimensionless noise amplitude for these two cell types? Before trying to address this question, let us first consider the third cell type discussed in this work, i.e. REF-52 [26]. In Fig. 3b, we present *p*(*θ*) for REF-52, focusing on experimental conditions similar to those used in the experiments presented in Fig. 3a, characterized by *ϵ* = 0.104 and *r* = 0.46. The latter two, together with the location of the maximum of *p*(*θ*) in Fig. 3b and in view of Eq. (5), imply *b* = 1.15, which is of course perfectly consistent with previous results [26]. Using then Eqs. (2) and (4) with a single fitting parameter 𝒯_eff_, we again obtain excellent agreement with the experimental data (solid line in Fig. 3b). The extracted dimensionless noise amplitude takes the value 𝒯_eff_ = 3.2*×* 10^*−*5^, which is of the same order of the one extracted for human aortic endothelial cells (cf. Fig. 3a), but nearly two orders of magnitude smaller than that of primary human umbilical cord fibroblasts.

To understand the origin of this difference, recall that 𝒯_eff_ = *τ D* and that the intrinsic cellular timescale *τ*, which corresponds to active remodelling of actin structures and focal adhesions, has been accurately determined only for REF-52 [26], but not for the two other cell types. However, we already know — as discussed above — that *τ* for primary human umbilical cord fibroblasts is more than an order of magnitude larger than that of REF-52. Consequently, most of the corresponding difference in 𝒯_eff_ = *τ D* is accounted for by the significantly larger *τ* of primary human umbilical cord fibroblasts, implying that their rotational diffusion coefficient *D* is in fact similar to that of REF-52. The interesting question of why these two cell types feature similar *b* and *D* values, but markedly different *τ* values, should be addressed in future work.

Whenever the value of *τ* is determined, most robustly by dynamic reorientation measurements and analysis (as done for REF-52 in [26]), the rotational diffusion coefficient can be obtained through *D* = 𝒯_eff_ */τ*. The rotational diffusion coefficient *D* in Eq. (3) can be independently measured by removing the external driving force after complete reorientation has been achieved and tracking the relaxation of an ensemble of cells back to a random orientational distribution *p*(*θ*) = 1*/π*. In particular, considering the order parameter *φ* = ⟨cos(2*θ*)⟩ (where the angle brackets denote ensemble average), the relaxation of an ordered orientational state characterized by *φ*_0_ *>* 0 takes the form *φ*(*t*) = *φ*_0_ exp(−*t/τ*_rel_), where the diffusional relaxation time is 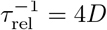 [35]. Applying this expression to REF-52, whose stationary orientational distribution under periodic driving is shown in Fig. 3b, we predict *τ*_rel_ =(4*D*)^*−*1^ = *τ* × (4 _eff_)^*−*1^ = 6.6 × (4×3.2 × 10^*−*5^)^*−*1^ sec ≃ 5 × 10^4^ sec. In fact, such an orientational relaxation experiment for REF-52 has already been performed in [17], and the diffusional relaxation time *τ*_rel_ that can be read off the up-pointing blue triangles in Fig. 3A therein is quantitatively consistent with our prediction. Very recently, the same procedure has been successfully applied to Caco-2 cells in a confluent epithelial layer [35].

It is important to highlight the qualitative difference between the rotational diffusion of cells and the rotational diffusion of molecules in thermal equilibrium. In the latter case, the rotational diffusion coefficient is proportional to the ordinary temperature. Cells, however, are much larger objects whose fluctuations at the cell scale are not determined by the ordinary temperature. Consequently, the rotational diffusion coefficient of cells, *D*, is related to the nonequilibrium, active nature of cells. Likewise, the relation 𝒯_eff_ = *τ D* is a generalized nonequilibrium Stokes-Einstein relation, where *τ* plays the role of a viscous-friction coefficient in the molecular, ther𝒯_eff_ mal equilibrium case. Indeed, recall that 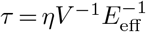, where *η* is an effective orientational viscous-like parameter. Clearly, these relations further justify the identification of 𝒯_eff_ as a dimensionless effective temperature, which naturally appears in the Boltzmann-like distribution in Eq. (4).

### INTRA-CELLULAR NEMATIC ORDER

Up to now we focused on cell body orientation under cyclic driving forces. This cell scale orientational state is accompanied by intra-cellular cytoskeletal organization that exhibits internal orientational order. Most notably, the cytoskeletal structure is characterised by discrete actin SFs whose orientation can be measured directly (cf. Figs. 4-6, to be discussed below). The level of internal orientational order is termed intra-cellular nematic order, inspired by the physics of nematic liquid crystals [44]. Intra-cellular nematic order, which provides a measure of the cytoskeletal polarization of cells, is relevant for various biological processes such as stem cell differentiation [45] and the self-organization of epithelial tissues [46]. Its dependence on the substrate’s rigidity in non-driven situations has been previously studied [36, 45]. Much less attention, however, has been given to the dependence of intra-cellular nematic order on periodic driving forces (see some discussion in the theoretical study of [32]). Our goal here is to study intra-cellular nematic order over a broad range of periodic driving conditions.

**FIG. 4.**
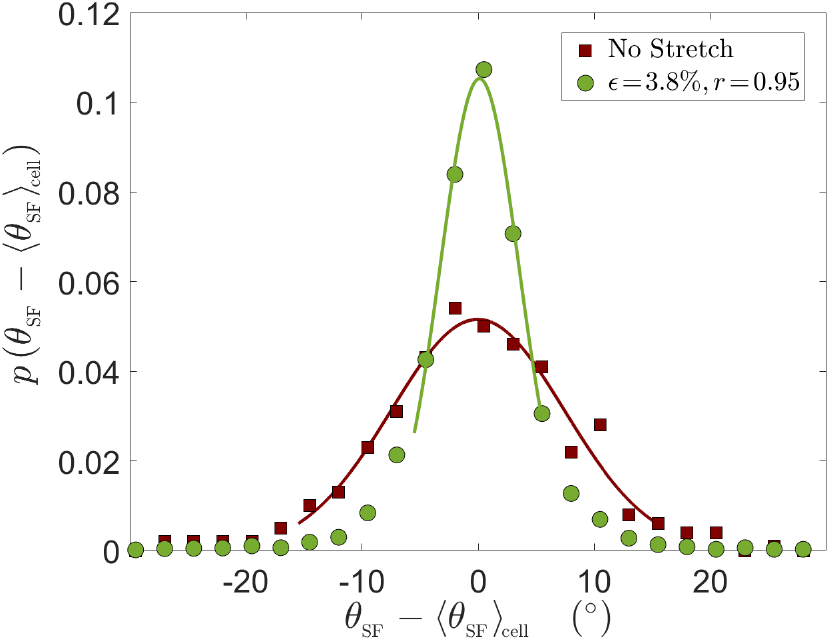
The stationary probability distribution function *p*(*θ*_SF_ *−* ⟨*θ*_SF_⟩_cell_) of the orientation of individual SFs within REF-52, where ⟨*θ*_SF_⟩_cell_ is the average stress fiber orientation of a given cell, in the absence of external driving (brown squares) and under periodic driving forces (green circles, see legend for the parameters). The solid lines are Gaussian fits to the central part of each distribution.

To that aim, we define *θ*_SF_ as the steady state (longtime limit) orientation of individual SFs within a cell. Next, we are interested in the probability distribution function of *θ*_SF_ *−*⟨*θ*_SF_⟩_cell_, where ⟨*θ*_SF_⟩_cell_ is the average stress fiber orientation of a given cell, shown in [26] to fully coincide with the cell body orientation (cf. Fig. 1b therein). The probability distribution function *p*(*θ*_SF_ *−*⟨*θ*_SF_⟩_cell_) allows one to quantify fluctuations in intra-cellular orientations. In Fig. 4, we first present *p*(*θ*_SF_*−* ⟨*θ*_SF_⟩_cell_) for REF-52 in the absence of external driving forces (brown squares), which serves as a reference case. We then superimpose on it the corresponding distribution obtained under the smallest strain used in our high-frequency cyclic stretching experiments (green circles), *ϵ* = 0.038. Both datasets have been obtained by analyzing ~ 80 cells and a few tens of SFs per cell, resulting in a few thousands of SFs in total. It is observed that the periodic driving results in reduced intra-cellular orientational fluctuations (a narrower distribution) compared to the reference case.

In order to quantify intra-cellular orientational fluctuations, we estimate for each experimental condition the distribution width *σ*_*θ*_ in the Gaussian approximation, obtained by fitting the central part of each distribution to a Gaussian, as shown in the solid lines superimposed on the experimental data in Fig. 4 (the tails of the distributions, which are generally non-Gaussian, are not discussed here). We use *σ*_*θ*_ to quantify intracellular nematic order. That is, a low value of *σ*_*θ*_ corresponds to a high level of intra-cellular nematic order, and vice versa. We extract *σ*_*θ*_, using the described procedure, in experiments employing various periodic driving conditions (*ϵ, r*), where the strain *ϵ* was varied in the range 0.038 *−* 0.104 and the biaxiality ratio *r* in the range 0.15 *−* 0.95 [26].

The resulting Gaussian widths *σ*_*θ*_ (*ϵ, r*) are presented in Fig. 5 (the error bars correspond to the uncertainty in each Gaussian fit), where *σ*_*θ*_ (*ϵ, r*) multiplied by *ϵ* is plotted against *r* (each value of *ϵ* is marked by a different symbol, see legend, and the non-multiplied data are shown in the inset). Our data span a factor of ∼3 in *ϵ* (from 3.8% to 10.4%, where for the smallest value we have only one *r* value), hence we cannot conclusively resolve the scaling with *ϵ*. Yet, the obtained results are consistent with *σ*_*θ*_ (*ϵ, r*) being inversely proportional to *ϵ* for a fixed *r*, i.e. with *ϵ σ*_*θ*_ (*ϵ, r*) *approximately* forming a function, which decreases with increasing *r*. Consequently, our results indicate that the intra-cellular nematic order increases with both increasing *ϵ* and biaxiality ratio *r*, i.e. that *σ*_*θ*_ decreases with both.

**FIG. 5.**
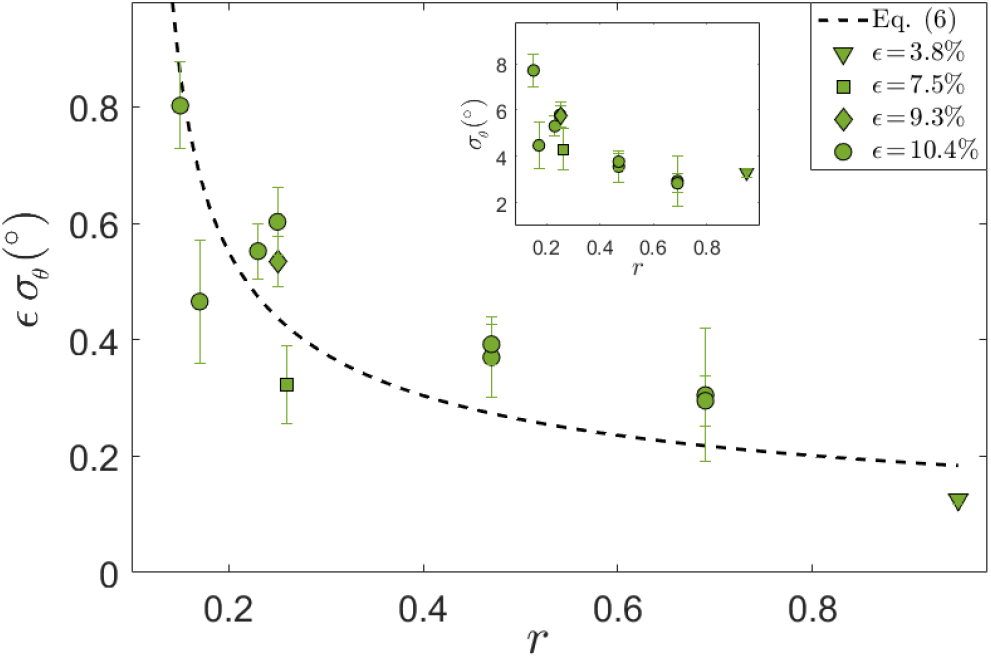
The Gaussian width *σ*_*θ*_ of *p*(*θ*_SF_ − ⟨*θ*_SF_⟩_cell_) for REF52 multiplied by *ϵ* vs. *r*, for different *ϵ* values (see different symbols and legend). The dashed line corresponds to Eq. (6) with 𝒯_eff_ = 3.9 × 10^*−*5^, see text for discussion. Note that in plotting the theoretical prediction in Eq. (6), angles have been converted from radians to degrees. (inset) The same as the main panel, but without multiplying *σ*_*θ*_ by *ϵ*.

How can one understand these trends? At the qualitative level, the fact that *σ*_*θ*_ (*ϵE, r*) decreases with increasing *ϵ*, and the fact that high-frequency periodic driving results in smaller *σ*_*θ*_ compared to the non-driven case (cf. Fig. 4), are expected due to the energetic penalizing effect of being oriented away from the minimum of the time-averaged elastic energy stored in the SFs, similarly to the biophysics implied by the cell body orientation theory in Eqs. (2)-(4). Moreover, a scaling relation *σ* (*ϵ, r*) *ϵ*^*−*1^ appears to be consistent with the quadratic energy functional *ū*(*θ*) of whole cells in Eq. (2). Yet, the theory in Eqs. (2)-(4) has been developed for cell body orientation at a coarse-grained level, treating the whole cell as a “composite material” composed of many SFs and their interconnections (as well as focal adhesions) [26], and as such cannot immediately apply to individual SFs.

At the same time, it is difficult to imagine that adjacent SFs reorient independently of their neighboring SFs, as there is a direct biophysical coupling between them [26, 42], and as they must experience some excluded-volume and nematic interactions. Consequently, one expects finite domains (or clusters) of adjacent SFs to orient in a correlated manner. In fact, in [26] it has been demonstrated that subcellular parts of a single cell can orient in *two different* mirror-image orientations, see Fig. 1b of the Supplementary Information file therein. Furthermore, it has been stated there in this context that: “This result indicates that the reorientation is not driven at the cell level, but rather involves a smaller part of it… and can be explained by SF clusters reorienting independently…”. Building on these ideas and observations, we speculate that the cell body theory of Eqs. (2)-(4) can be used as a rough approximation for *p*(*θ*_SF_ *−*⟨*θ*_SF_⟩_cell_), and in particular for *σ*_*θ*_ (*ϵ, r*).

To further substantiate this suggestion, we present in Fig. 6 an image of a single REF-52 under periodic driving (*ϵE* = 0.104 and *r* = 0.17) in the long-time limit, where SFs were imaged after being stained with fluorescently labelled phalloidin. We also added as a guide to the eye (dashed line) the cell body orientation predicted theoretically in Eq. (5). It is observed that domains composed of several SFs are locally oriented in a correlated manner in a direction that does not necessarily coincide with the cell body direction (some of them are marked by ellipses that serve as guides to the eye). This observation qualitatively supports the suggestion stated above (note that no attempt is made here to quantify the correlated SF domains, e.g. their correlation length).

**FIG. 6.**
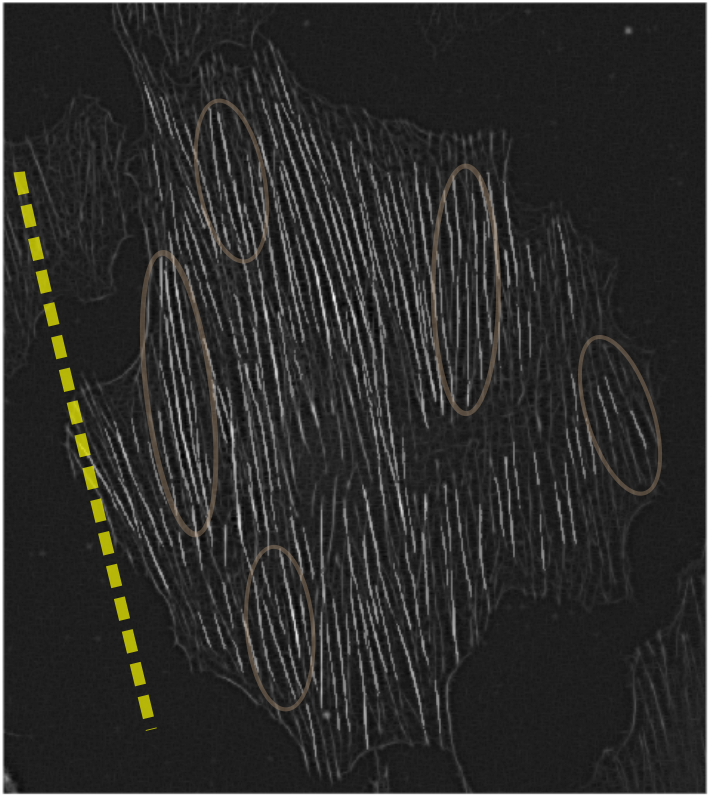
An image of a single REF-52 under periodic driving (*ϵ* = 0.104 and *r* = 0.17) in the long-time limit, where SFs were imaged after being stained with fluorescently labelled phalloidin. The dashed line corresponds to the cell body orientation predicted theoretically in Eq. (5), added as a guide to the eye. The ellipses, also added as guides to the eye, illustrate the possible existence of correlated SF domains, see text for additional discussion.

With this evidence in mind, we assume now that Eqs. (2)-(4) also apply to SF domains and hence can teach us something about intra-cellular orientational fluctuations. We can then use Eq. (2) inside Eq. (4), and derive the Gaussian approximation around the peak of the distribution at *θ*_m_. The resulting Gaussian width *σ*_*θ*_ (*ϵ, r*) takes the form

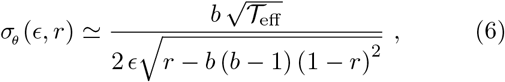

which is trivially proportional to *ϵ*^*−*1^. As *b* = 1.13 for REF-52 has been determined by the mean-field reorientation dynamics [26], Eq. (6) contains a single unknown parameter, the dimensionless noise amplitude𝒯_eff_. In fact, pushing the analogy between SF domains and whole cells a bit further, we expect them to experience similar levels of dimensionless orientational noise, and hence expect 𝒯_eff_ for SFs to be close to 3.2 × 10^*−*5^, the value obtained in Fig. 3b for whole cells. Indeed, upon superimposing (dashed line) the prediction in Eq. (6), with𝒯_eff_ = 3.9*×* 10^*−*5^, on top of the experimental data in Fig. 5, a reasonable agreement is revealed. Consequently, while applying the cell body fluctuations theory to SF domains is clearly approximate and somewhat rough, Eq. (6) appears to semi-quantitatively capture the experimental trends in intra-cellular nematic order as a function of the driving parameters *ϵ* and *r*.

## SUMMARY AND OUTLOOK

In this work, we first experimentally validated two theoretical predictions regarding the timescale and stiffness scale separation associated with the reorientation of adherent cells in response to periodic driving. In particular, we showed that cellular reorientation occurs when the driving frequency *f* is much larger than an inverse intrinsic cellular timescale *τ* ^*−*1^ and when the substrate stiffness *E*_sub_ is much larger than the (high-frequency) effective cell stiffness *E*_eff_, as predicted. As such, these results further support an established single-cell mean-field theory of cellular reorientation under periodic driving [26]. With this fundamental mean-field level understanding of cellular reorientation at hand, we turned our focus to the stationary orientational distribution function *p*(*θ*).

We extended the mean-field theory to include uncorrelated, additive active noise and compared the resulting stationary probability distribution functions of cellular orientation to extensive experimental data for three different cell types. These theoretical predictions quantitatively agree with the experimental data for cell body orientation, and allowed to extract various cellular parameters, most notably the dimensionless measure of cellular elastic anisotropy *b* — which is determined by the most probable orientation — and the dimensionless noise amplitude 𝒯_eff_. We found that cellular elastic anisotropy is almost constant, attaining values in the range *b* = 1.12*−*1.23, across the three cell types considered here. The dimensionless noise amplitude 𝒯_eff_ was found to be significantly larger for one cell type, a difference that was attributed to the significantly larger intrinsic timescale *τ* of this cell type.

Once the intrinsic timescale *τ* is known, typically through dynamic (time dependent) reorientation measurements of *θ*(*t*) in the *f* »*τ* ^*−*1^ limit [26], the cellular rotational diffusion coefficient is obtained from *D* = 𝒯_eff_ */τ* The latter, as explained above, is intimately linked to the nonequilibrium, active nature of cells and provides a quantitative prediction for the free relaxational dynamics of initially ordered cells upon which orientational order is lost, i.e. when the periodic driving forces are removed and a random distribution *p*(*θ*) = 1*/π* is approached. From a broader perspective, our work provides concrete guidelines for how the widespread cyclic stretching experimental framework — where *E*_sub_, *ϵ, r* and *f* are controlled — can be systematically used for extracting the basic cellular parameters *b, τ* and *D*, which provide insight into the mechanical properties of cells and what governs them. To this end, the design of various experiments and the biophysical observables accessible through them are summarised in a flowchart available in Appendix C.

Finally, we considered the effect of periodic driving on intra-cellular nematic order as manifested in cytoskeletal organization, in particular in the orientation of actin SFs. We found that intra-cellular nematic order increases with both the magnitude *ϵ* of the periodic driving forces and the biaxiality strain ratio *r*. These results were semi-quantitatively explained by applying the same cell body fluctuations theory to orientationally correlated SF domains. Overall, in this work we offer a quantitative understanding of active orientational fluctuations in living cells under period driving, both cellular and intracellular, which are relevant for physiological conditions.

Our results also raise several questions and open future investigation directions. It would be very interesting to extract the cellular properties *b, τ* and *D* for many cell types, and to explore the biophysical factors and molecular processes that control them. In particular, it would be interesting to explore the possible relations between temporal cellular parameters *τ* and *D*, and the kinetics of adhesion proteins and actin turnover rates [47, 48]. In addition, it would be interesting to explore the possible relations between the rotational diffusion coefficient *D* and cell migration. Finally, it would be interesting to further explore the quasi-universality of the level of cellular elastic anisotropy (in the high-frequency limit), as quantified by *b* = 1.12*−* 1.23. It may suggest generic cytoskeletal organization principles across different cell types, which should be better understood. It is interesting to note in this context that for Caco-2 cells in a confluent epithelial layer, a larger value of *b* = 2.25 has been estimated [35], apparently due to cell-cell interactions.

The structure of Eqs. (2) and (4) shows that the characteristic width of *p*(*θ*) scales as 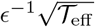. It implies that a population of cells will exhibit enhanced orientational correlations with increasing*ϵ*under periodic driving. Likewise, the cytoskeletal organization of cells features enhanced nematic order, i.e. cell polarization, with both increasing *ϵ* and *r*. It would be interesting to explore whether these principles are at play in physiological processes, e.g. during tissue development where well-coordinated inter-cellular organization is necessary. Finally, in relation to applications, these biophysical principles can be used in the context of tissue engineering — e.g. for cardiovascular regeneration — where controlled cell-to-cell alignment is a major challenge of prime importance [8, 9, 11].

## Acknowledgements

E.B. acknowledges support from the Ben May Center for Chemical Theory and Computation and the Harold Perlman Family.

## APPENDIX

### A. Cellular orientation experiments and data analysis

The cellular orientation data of primary human umbilical cord fibroblasts, presented and analyzed in Fig. 2, correspond to the experiments of [18]. The cellular orientation data of human aortic endothelial cells, presented and analyzed in Fig. 3a, correspond to the experiments of [19]. The histograms originally reported on in these two papers [18, 19] were digitized directly from figures therein (see also the captions of Fig. 2-3). In cases in which the reported quantity was the probability *P* (*θ*) (rather than the probability distribution function, such that ∑ *P* (*θ*) = 1), we normalized the latter by the average separation *δθ* between consecutive probability entries/bin values in order to obtain the probability distribution function *p*(*θ*) *≡ P* (*θ*)*/(δθ)*. The latter satisfies *p*(*θ*)*dθ ≈ ∑ p*(*θ*) ⟨ *δθ*⟩ = 1. The cellular orientation data of REF-52, presented and analyzed in Fig. 1 and Fig. 3b, correspond to the experiments some of us reported on in [26], see experimental details therein.

In the context of experimentally extracting the orientational distribution *p*(*θ*), we would like to raise here a methodological issue, which has not received enough attention in the literature. The distributions for primary human umbilical cord fibroblasts presented in Fig. 2 were obtained using ensembles ranging from 397 to 1493 cells for each experimental condition [18], presumably leading to proper statistical convergence. On the other hand, the distribution for human aortic endothelial cells presented in Fig. 3a was obtained using 77 *−*144 cells (the exact number is not specified in [19]) and the distribution for REF-52 presented in Fig. 3b was obtained using 76 cells (our own data, which as stressed above, were originally obtained for studying single-cell reorientation dynamics [26]). It is desired in future work to employ larger cell ensembles to ensure the statistical convergence of experimental data and to minimize potential statistical undersampling effects in *p*(*θ*).

To test the theoretical prediction in Eq. (4), together with Eq. (2), against the experimental data for the three cell types, we employed a fitting procedure to be explained next (and whose outcome is plotted in the solid lines in Figs. 2-3). The fitting procedure employed MATLAB’s [49] fmincon function, which allows to find the minimum of constrained nonlinear multivariable functions. For the latter, we defined a standard quadratic error function between the experimental distributions and *p*(*θ*) of Eq. (4), in the form err *≡* [*p*(*θ*_M_ ; *r, b, 𝒯*_eff_) *− p*_M_]^2^, where *θ*_M_ and *p*_M_ are the measured orientations and probabilities, respectively. The main fitting parameter is 𝒯_eff_, while *r* and *b* are either highly constrained or fixed.

The value of *r* is, in principle, determined by the experimentalist. However, its actual value depends on the precise loading configuration employed in a given experiment [26], and the mapping between the latter and *r* is not generally known, and hence *r* should be measured. When such accurate measurements are available, as is the case the experiments whose data appear in Fig. 3, *r* is just fixed. In the experiments whose data appear in Fig. 2, the *r* values have been measured and reported on with some uncertainty. In this case, we allowed for variations within the range *r* = *r*_M_ *±* Δ*r*_M_, with *r*_M_ = 0.15 and Δ*r*_M_ = 0.05 for Figs. 2a-e, and *r*_M_ = 0.29 and Δ*r*_M_ = 0.05 for Figs. 2f (the uncertainty level of Δ*r*_M_ = 0.05 is the one reported in [18]).

The *b* values were determined based on the most probable orientation, as described next. We fitted 5 data points in the vicinity of the most probable data point (i.e. 2 points to the left of it and 2 to the right) to a quadratic function *Aθ*^2^ + *Bθ* + *C*, such that the most probable orientation is obtained as *θ*_m_ = *− B/*2*A* (which does not necessarily coincides with the most probable data point). When the parabolic fit fails, we used the most probable data point as an estimate of *θ*_m_. Then, the prediction in Eq. (5), together with the value of *r*, were used to estimate *b*, which in turn was used as an initial value in the constrained nonlinear minimization. Finally, in order to minimize the possible sensitivity of the fitting process (i.e. the constrained minimization of the error function) to the initial values of the fitting parameters, we considered an ensemble of 200 random initial values for *r* and *b* (within their respective constrained ranges), and for 𝒯_eff_ for each case. The reported results (solid lines in Figs. 2-3) correspond to the smallest error within this ensemble.

### B. Intra-cellular nematic order data (stress fibers orientation analysis for REF-52)

The intra-cellular nematic order data for REF-52, appearing in Figs. 4-6, correspond to the stress fibers (SFs) orientation analysis of the experiments some of us reported on in [26]. For each cell (under period driving, in the long-time limit), SFs have been detected (following the image analysis detailed in [26]) and their individual orientation *θ*_SF_ has been measured. As stated in the main text, there are a few thousands SFs per experimental condition, hence the resulting probability distribution function *p*(*θ*_SF_ *−* ⟨ *θ*_SF_ ⟩_cell_) (which includes data from *∼*7080 cells and ⟨*θ*_SF_⟩_cell_ is the average per cell) is statistically converged (e.g. largely insensitive to the binning scheme employed). Two examples of *p*(*θ*_SF_ *−* ⟨*θ*_SF_ ⟩_cell_) are presented in Fig. 4.

The central part of each resulting distribution was found to be predominantly Gaussian. Hence, we fitted the central part of each distribution (including data points extending at least to half of the maximum of the distribution) to a Gaussian and extracted its standard deviation *σ*_*θ*_. The uncertainty in the values of *σ*_*θ*_, reflected in the error bars in Fig. 5, correspond to the associated uncertainty in the Gaussian fit. The theoretical curve in Fig. 5 (dashed line therein) corresponds to a single-parameter fit of Eq. (6) to the experimental data (with 𝒯_eff_ being the only fitting parameter), using the same constrained nonlinear minimization procedure discussed above.

### C. Design of experiments and biophysical observables

Here we briefly summarise the design of various experiments and the biophysical observables accessible through them, as explained in this work. This is done in the flowchart presented in Fig. 7, see also the figure caption.

**FIG. 7.**
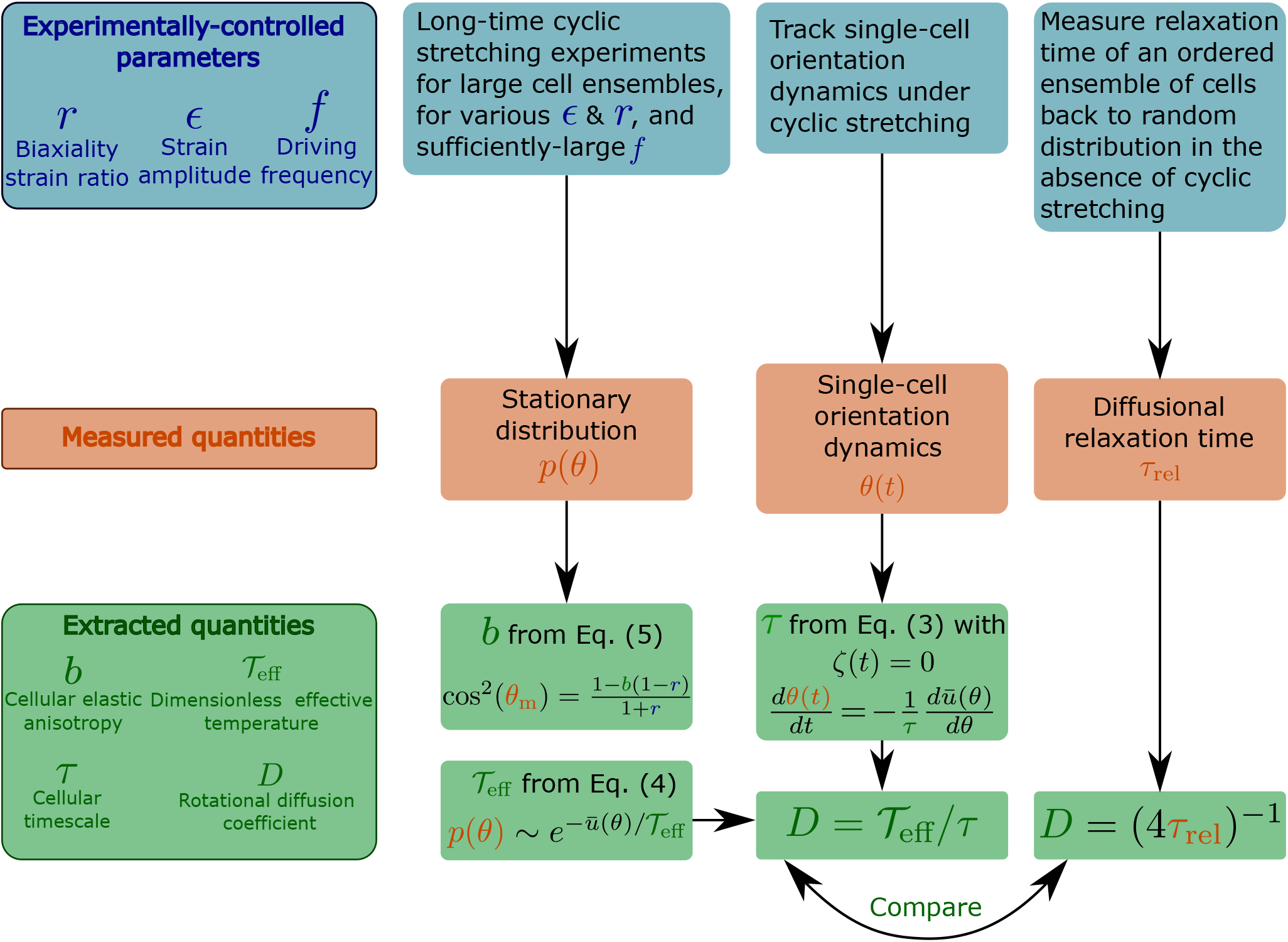
A flowchart that briefly summarises various cyclic stretching experimental setups, the measured observables and the subsequent extracted properties of cells. The experiments, represented by the blue boxes (top row), control the driving force (cyclic stretching) parameters *r* (biaxiality strain ratio), *ϵ* (strain amplitude) and *f* (driving frequency). The elastic properties of the substrate to which the cyclic stretching is applied, e.g. its stiffness *E*_sub_ (not marked on the flowchart), are also controlled. Note that the biaxiality ratio *r* generally depends on both the experimental loading configuration (e.g. the stretcher/stretchers employed, its/their coupling to the substrate etc.) and the substrate’s elastic properties, and hence should be extracted from direct measurements in the cells’ vicinity. The measured observables appear in the orange boxes (middle row) and include: (i) the cellular orientation probability distribution function *p*(*θ*), obtained by long-time cyclic stretching experiments performed on a large ensemble of non-interacting cells (the ensemble size should be selected so as to ensure statistical convergence, which should be quantitatively tested and demonstrated), (ii) the reorientation dynamics *θ*(*t*) of single cells from their original random orientation to one of the mirror-image steady state orientations. It requires tracking single cells over a long period of time with sufficient spatiotemporal resolution, (iii) the diffusional relaxation time *τ*_rel_, obtained by tracking the relaxation dynamics of an ensemble of cells from their cyclic stretching induced ordered state to a random orientational state upon the removal of the driving forces. These observables allow the extraction of the basic cellular properties *b*, 𝒯_eff_, *τ* and *D*, appearing in the green boxes (bottom row). *b* is a dimensionless measure of cellular elastic anisotropy (in the high-frequency limit), which is intimately related to the cytoskeletal organization of cells and their intrinsic polarization. It is extracted from the position of the maximum of *p*(*θ*), denoted by *θ*_m_, and the biaxiality ratio *r* using Eq. (5). Note that the latter equation is valid for 1 *−*1*/b*≤ *r* ≤1*/*(*b* − 1), which is very common. For 0 *< r <* 1 *−*1*/b*, we have *θ*_m_ = 90^*°*^, and *b* is determined as a fitting parameter when *p*(*θ*) is fitted to Eq. (4) (with *ū*(*θ*) of Eq. (2)). This fitting procedure allows to extract the dimensionless effective temperature 𝒯_eff_, which provides a dimensionless measure of cellular orientation fluctuations. Since the latter satisfies a generalized Stokes-Einstein relation of the form 𝒯_eff_ = *τ D*, the rotational diffusion coefficient of cells *D* can be obtained once *τ* is extracted. *τ* is an intrinsic remodeling cellular timescale, which is related to cells’ ability to disassemble and reassemble their adhesion complexes and actin structures during driven reorientation dynamics. It can be obtained from the single cell dynamics *θ*(*t*) using Eq. (3), once the noise term is removed. The rotational diffusion coefficient of cells *D*, which is intrinsically linked to the nonequilibrium active nature of cells, is determined through *D* = 𝒯_eff_ */τ* and also independently through the measurement of the diffusional relaxation time *τ*_rel_. The outcome of the two independent procedures should be compared.

